# Transcriptome analysis and connectivity mapping of *Cissampelos pareira* L. provides molecular links of ESR1 modulation to viral inhibition

**DOI:** 10.1101/2021.02.17.431579

**Authors:** Madiha Haider, Dhwani Dholakia, Aleksha Panwar, Parth Garg, Atish Gheware, Dayanidhi Singh, Khushboo Singhal, Shaunak A Burse, Surekha Kumari, Anmol, Arjun Ray, Guruprasad R. Medigeshi, Upendra Sharma, Bhavana Prasher, Mitali Mukerji

## Abstract

Bioactive fractions or compounds obtained from medicinal plants have been used for the treatment of multiple diseases. This effect could be due to common pathways underlying these conditions that are targeted by such medicines. In this study, we explored the molecular basis of action of one such herbal formulation *Cissampelos pareira*, used for the treatment of female hormone disorders and fever. Genome-wide expression studies on MCF7 cell lines treated with Cipa extract were carried out using Affymetrix arrays. Transcriptome analysis revealed a downregulation of signatures of estrogen response governed by estrogen receptor α (ERα). Molecular docking analysis identified 38 constituent molecules in Cipa that potentially bind (ΔG< -7.5) with ERα at the same site as estrogen. Cipa transcriptome signatures show high positive connectivity (https://clue.io/) scores with protein translation inhibitors such as emetine (score: 99.61) and knockdown signatures of genes linked to the antiviral response such as ribosomal protein RPL7 (score: 99.92), which is also an ERα coactivator. Cipa exhibits antiviral activity in dengue infected MCF7 cells that is decreased upon ESR1 (estrogen receptor 1) gene knockdown. This approach reveals a novel pathway involving ESR1-RPL7 axis that could be a potential target in dengue viral infection.

## Introduction

Plant-based formulations are often used to treat multiple disorders. The pharmacological effects that the drug substances produce may be due to the interaction with target molecules at the molecular level to induce a change in the target molecule functioning. Multiple active principles present in a single herb, could be postulated to target different pathways to bring about synergistic effects. It has been observed that therapeutics that have more than one target may possess in principle a safer profile compared to single-targeted ones. The use whole formulations instead of a herb’s active principles may bring about maximum efficacy and minimize drug resistance that is commonly set in with a single molecule ^1–3^. There, however, remains a gap in understanding the molecular underpinnings of whole formulations.

### Cissampelos pareira

L. (Cipa) a popular herbal medicine, has traditionally been used for fever, contraception and other reproductive system related disorders ^4–6^. It is also widely used for treating asthma, cough, skin ulcers, snakebite, and jaundice in many parts of the world ^5^. Alcoholic and aqueous alcoholic extracts of Cipa have been shown to have antiviral and antimalarial activities, respectively ^4,7^. Cipa methanolic extracts have also been shown to potentially modulate the estrus cycle in mice ^8^. However, there is limited research on exploring whether there are shared pathways through which these extracts work in such diverse diseases. We used a genomics framework to explore the molecular signatures of the whole formulation of Cipa and to probe whether there are molecular players that connect the hormonal axis with antiviral inhibition.

In this study, we carried out a transcriptomic analysis of Cipa in MCF7 cell lines and queried its involvement in different pathways using gene ontology tools, Gene Set Enrichment Analysis, and probed its connectivity with drugs and other perturbagens including gene knockdowns using L1000 ^9,10^ in CMAP (the connectivity map). Pathways linked to lipid metabolism, viral transcription, translation, and estrogen axis seem to be modulated by CIPA. It also shows a high connectivity score with protein synthesis inhibitors that have demonstrated antiviral effect in *in vitro* studies. Besides the downregulation of ESR1, we also obtained a high connectivity score of Cipa with a knockdown signature of a selective estrogen receptor-alpha coactivator, RPL7 also involved in protein translation. Docking studies of CIPA constituents to estrogen receptor alpha show a high binding affinity of ∼ 38 compounds to regions that bind known ERα modulators. CIPA inhibits dengue virus replication in MCF7 cell lines, and ESR1 knockdown in CIPA treated infected cells reduces this effect.

In trying to connect the dots between diverse activities of a commonly used herbal medicine, we have been able to identify an interesting link between hormone response and antiviral activity. This framework could be used investigate other similar medicines and identify novel and targetable links between diseases.

## Results

### Cipa modulates estrogen receptor alpha in MCF7 cells

In order to probe the mechanism of action of *Cissampelos pareira*, we performed transcription profiling on Affymetrix microarray HTA 2.0. Genome wide expression assay of MCF7 cells provided 93 downregulated and 131 upregulated genes at 500μg and 253 downregulated and 587 upregulated genes at 1000μg, of Cipa treatment (**Supplementary file 1**). Gene ontology analysis revealed that the upregulated set had significant enrichment of transcription regulation by RNA polymerase II, tissue development, supramolecular fiber organization etc., whereas the downregulated genes were enriched in lipid metabolism processes (**Fig S3**).

Gene Set Enrichment Analysis (GSEA) analysis provided 59 positively and 12 negatively enriched gene sets (p-value <0.05 and FDR< 5%). We observed a significant enrichment of estrogen response gene sets in both positive and negative enrichments (**Figure 1A**). ESR1 mRNA expression was observed to be downregulated in the array that was confirmed by real-time quantitative analysis of mRNA levels in both concentrations of Cipa (**Figure 1B**). Motif searches of estrogen response elements (ERE) reveal significant enrichments in ERE density in up and downregulated genes (p value= 0.0054) in the 5KB upstream region. There is no significant difference in the number of ESR1 elements between up regulated, downregulated and unchanged genes sets in the 5 KB upstream region (**Figure 1C**). The regression values between fold change and the number of ERE sites revealed the presence of ERE sites in the 5KB upstream region have negative effects on expression. The differentially regulated genes also include well-characterized estrogen response genes with EREs in their promoters (**Fig S4**).

**Figure 1.**
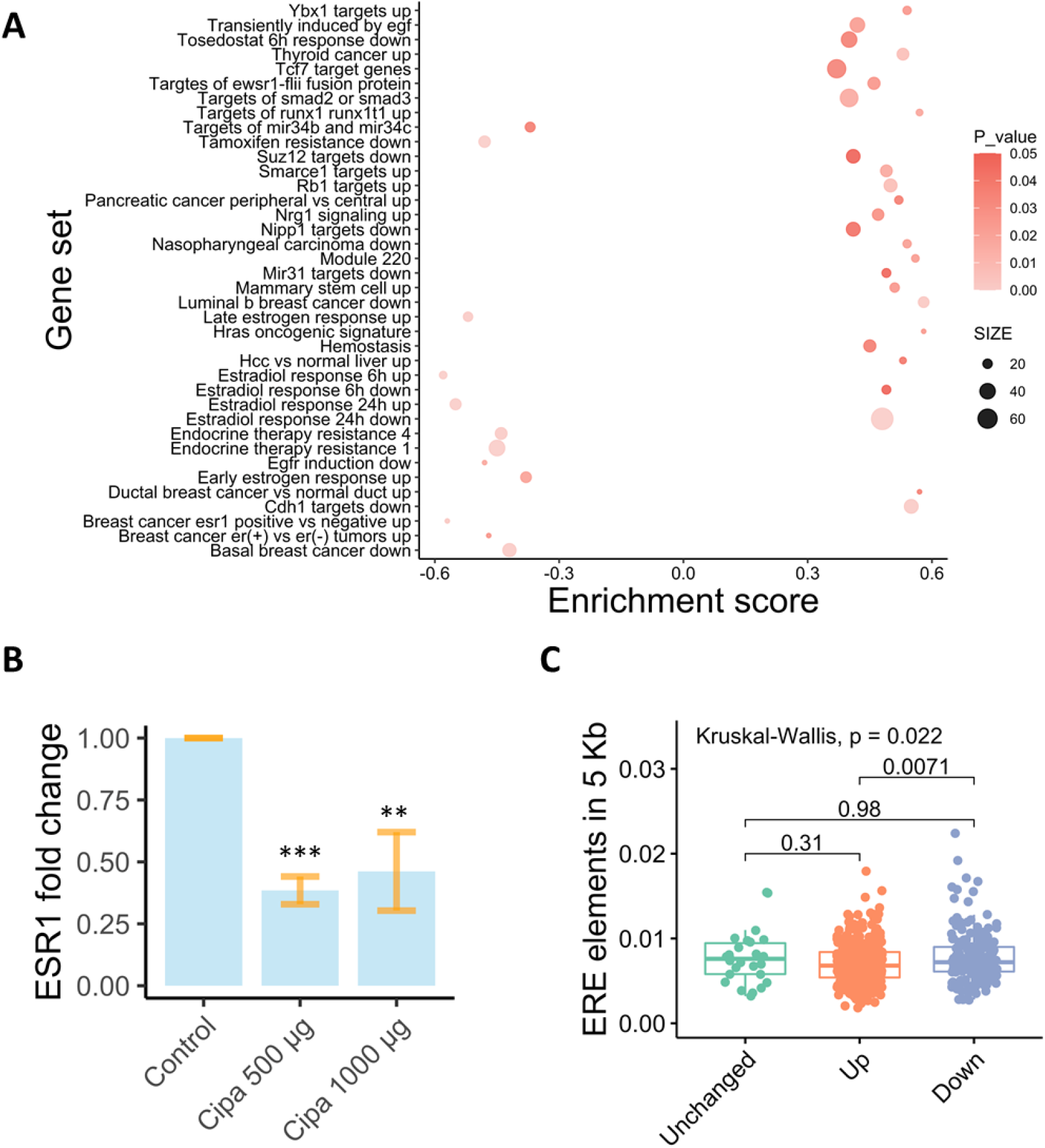
Estrogen receptor modulation by Cissampelos pareira. A) Gene sets enriched for genes differentially expressed, positive enrichment score for upregulated and negative score for downregulated genes. p-value <0.05 and FDR< 5%. B) Fold change in mRNA expression of ESR1 transcript at 500 and 1000µg. Graph is cumulative of three separate experiments (n=3, N=3). *** p-value<0.001, ** p-value<0.01. C) Density of estrogen response elements 5Kb upstream of differentially expressed genes, the ERE elements are higher in density in down regulated genes than upregulated genes (p value<0.05).

### CIPA shows high positive connectivity scores with protein synthesis inhibitors that are potential antivirals

A ranked gene signature of 69 down and 129 up genes was used to query the connectivity map (https://clue.io/). At a cut-off score of 95 there were 69 positively connected small compounds. The top-scoring compounds belonged to the class of protein translation inhibitors e.g., emetine (99.61), cycloheximide (99.86), verrucarin-a (99.47), cephaeline (99.51) (**Figure 2A**). Emetine, homoharringtonine, anisomycin, cephaeline, cycloheximide, and QL-XII-47 have reported antiviral activity (**Table S2**). In the genetic perturbations set, 77 genes had knockdown (KD) signatures score >90, and all were involved in translation initiation, and viral transcription (**Figure 2B, Fig S6**). Since protein synthesis inhibition appears to be significant in our CMAP analysis, we decided to explore our expression data for genes involved in translation. The mRNA levels of the gene with the highest KD connectivity, RPL7 was downregulated after Cipa treatment (**Figure 2C**). In vitro assays showed moderate inhibition of translation in response to Cipa (**Fig S7**).

**Figure 2.**
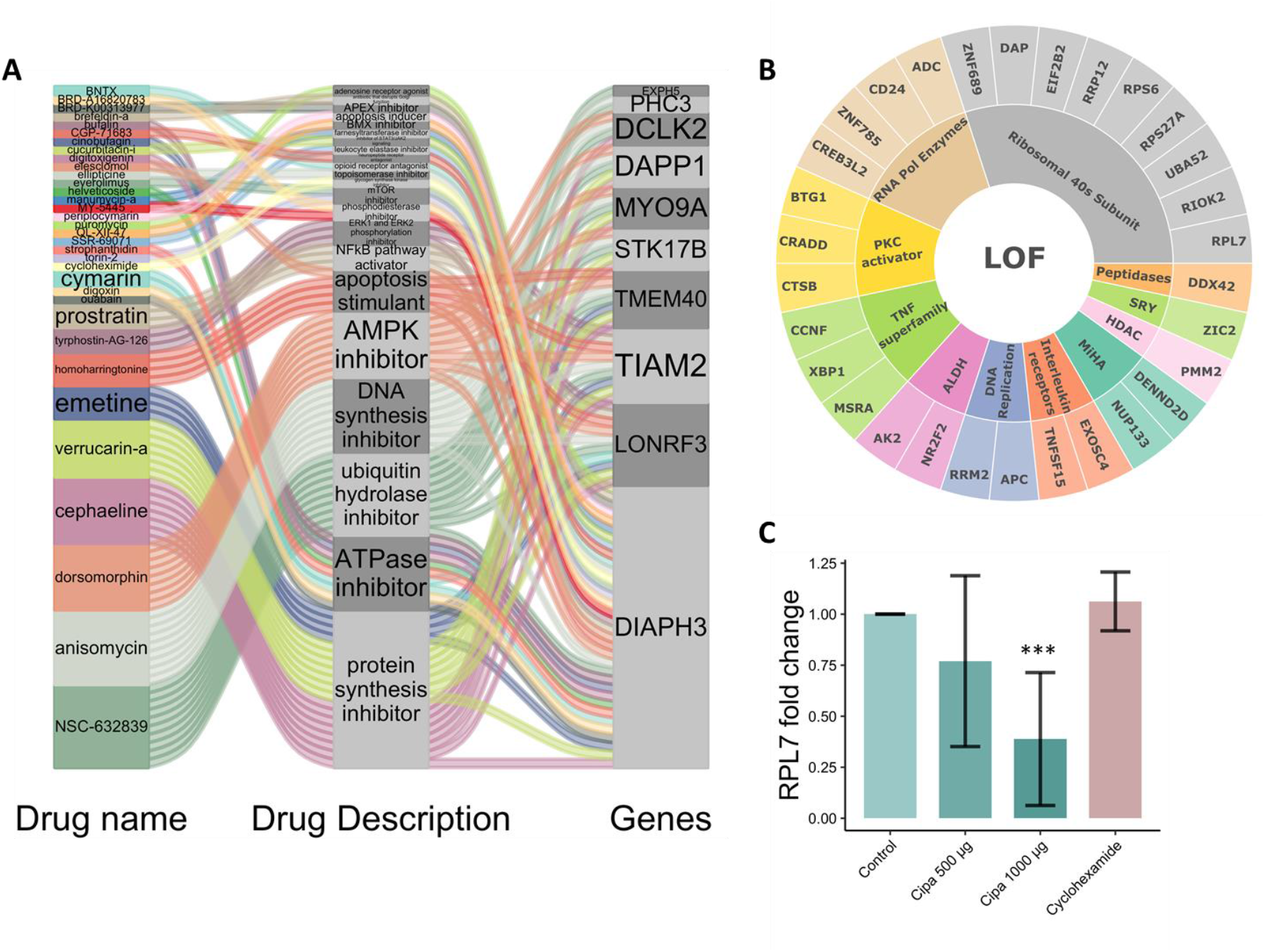
Connectivity map analysis of Cipa gene expression signature: A) Alluvial plot depicts small compounds with highest connectivity score >99, the class of perturbation they belong to such as protein synthesis inhibitor and the genes common with the signatures. B) Sunburst diagram shows the frequency of 11 perturbation types to which the top 30 loss of function signatures belong. C) mRNA expression of RPL7 gene in response to Cipa, Graph is cumulative of two separate experiments (n=2, N=3)*** p-value<0.001

### Virtual screening demonstrates Cipa constituents bind with Estrogen receptor alpha

To investigate the atomistic detail of possible Cipa constituent binding to the Estrogen receptor alpha, unbiased virtual screening of the constituent compounds against the receptor was employed. This revealed four preferential locations of binding for the 61 compounds. The largest cluster (pink) comprised 38 out of the 61 Cipa constituents, followed by the cluster of 8 (blue), 6 (orange), and 3 (green). As a positive control, we took previously reported SERMs, namely, tamoxifen and 17-*β*-estradiol. Our results show that both of these control molecules bind at the same location as the largest cluster in our findings with high concurrency to the reported binding energy (**Figure 3, Table 1**). It is also noteworthy that more than 30 Cipa components had a minimum binding of <-8.00 kJ/mol and at least 9 molecules have a minimum BE of <-9 which is equal to or even higher than Tamoxifen binding energy -9.2 (**Table S1**). Ligplot of the detailed binding sites for the controls and top binding molecules in **Fig S5**.

**Table 1:**
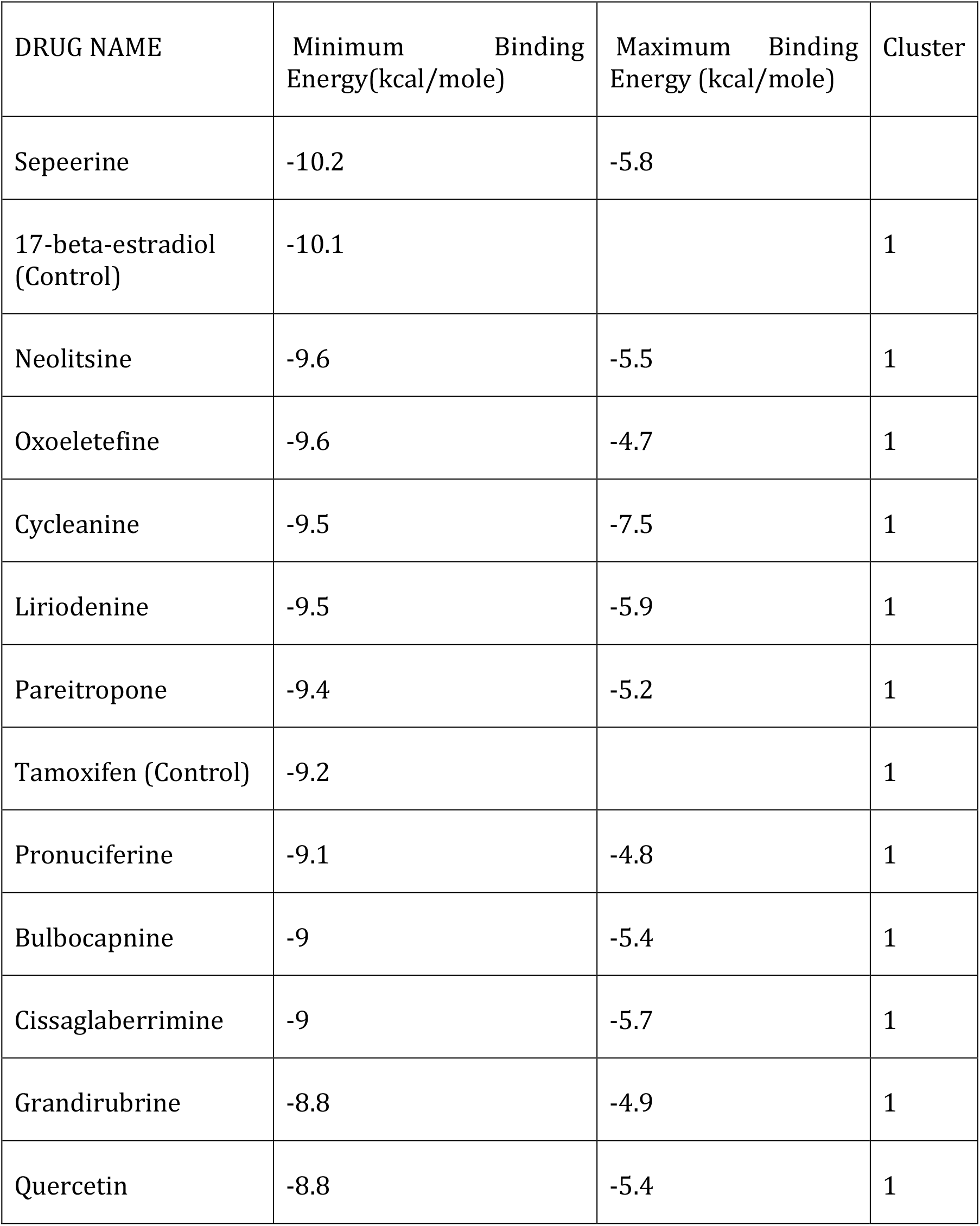
Cipa constituent compounds, their minimum and maximum binding energies with ER-alpha, the binding cluster each fall into (full table in supplementary)

**Figure 3.**
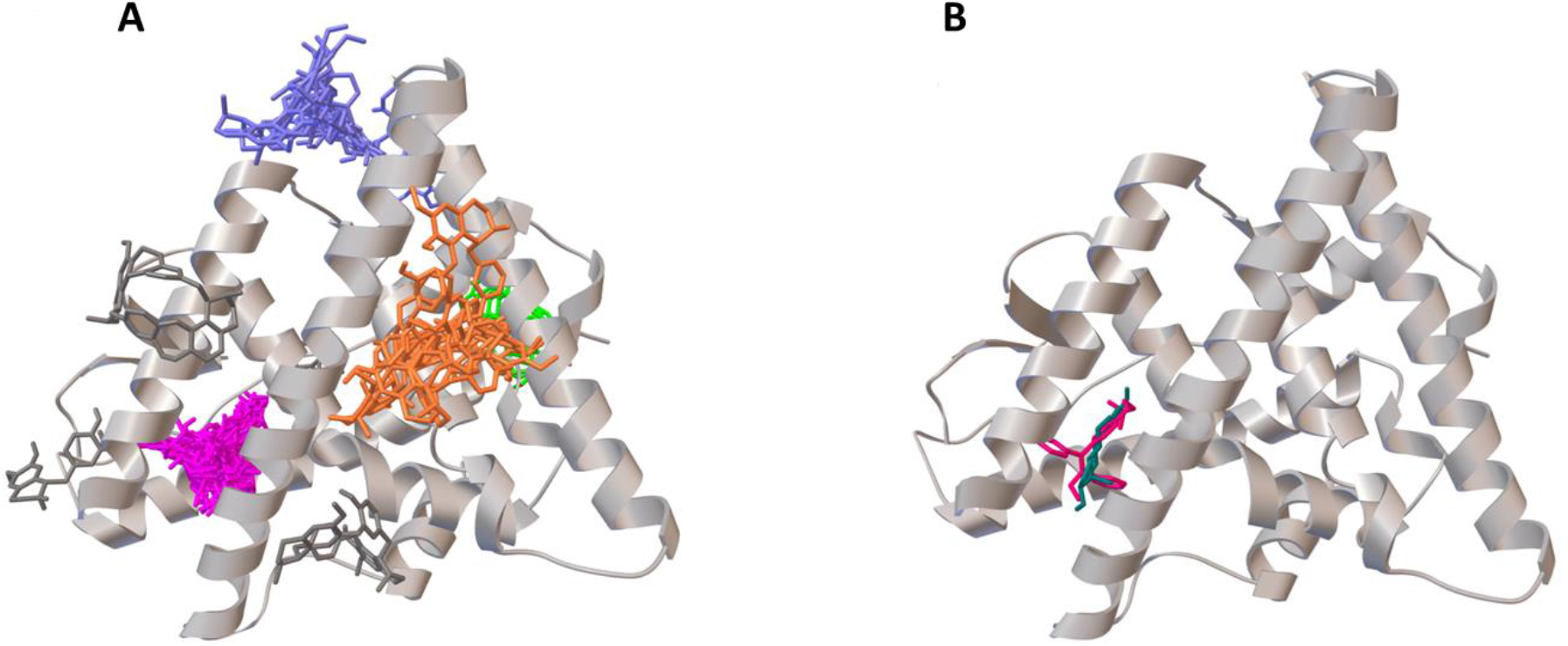
Cipa constituents can potentially bind with ERα at four distinct sites: A) Virtual docking sites of 61 Cipa constituents (four clusters, pink, green, orange and blue). B) Binding site of controls tamoxifen (pink) and 17 beta estradiol (blue) on estrogen receptor alpha.

### Cipa inhibits DENV infection in breast cancer cell lines in ESR1 dependent manner

To explore ESR1 involvement in antiviral activity of Cipa, we infected two cell lines that vary in estrogen receptor alpha expression, namely MDA MB 231 and MCF7 that have minimal and high expression of ESR1 respectively (**Figure 4A**) with DENV-2 (**Fig S9**). Both cell lines were infected with DENV-2 at 10 MOI followed by treatment with Cipa. We observe that dengue inhibition was higher in MCF7 cells compared to MDA-MB-231 cells at both concentrations of Cipa (**Figure 4B**). This differential inhibition could be a consequence of different cell types. To further validate that the DENV inhibition is ESR1-dependent, we performed siRNA-mediated knockdown of ESR1 in MCF7 cells. Knockdown efficiency as confirmed by qRT-PCR 48 h post transfection show approximately 70-80% reduction in mRNA levels compared to that of non-targeting control (NTC) siRNA. After viral infection and subsequent Cipa treatment we observe that the inhibitory effect of Cipa was diminished in ESR1-knockdown compared to mock-treated cells suggesting that DENV inhibition due to Cipa is ESR1-dependent in MCF7 cells. (**Figure 4C, D**).

**Figure 4.**
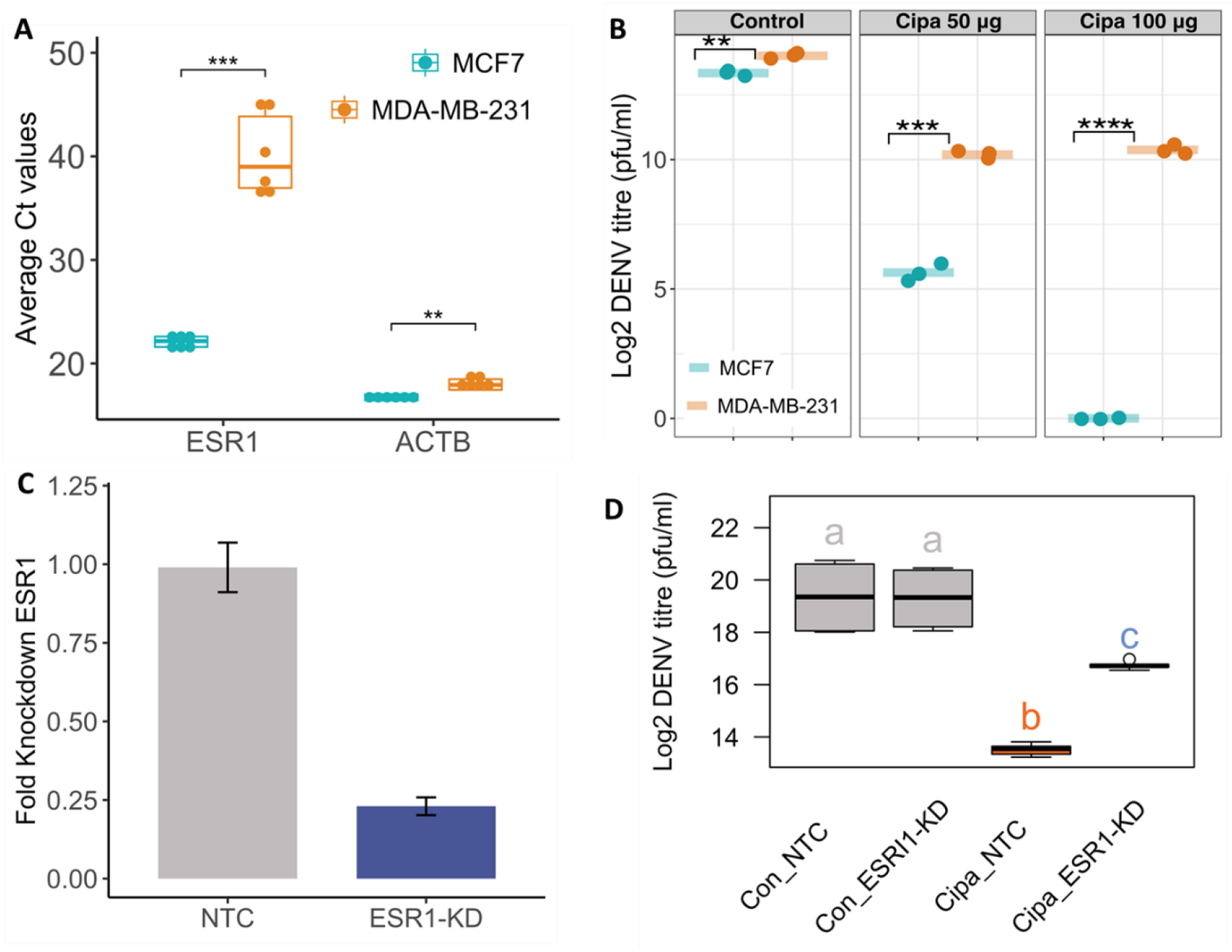
DENV inhibition by Cipa in ESR1-dependent manner: A) Average Ct value of ESR1 and ACTB in MCF7 and MDA-MB-231 cells. B) Both cell lines were infected with DENV at 10 MOI and after viral adsorption, Cipa was added at two different concentrations of 50 and 100μg. Viral titers in the supernatant were determined by plaque assay. Graph indicates DENV infection titers at 24 h pi. ** p-value<0.01 ***p-value<0.001 ****p-value<0.0001. C) MCF7 cells were treated with siRNA specific to *ESR1* and scrambled siRNA as a control (labeled as NTC (nontargeted control). Graph indicated knockdown efficiency determined at mRNA level by qRT-PCR at 48 h post transfection. D) At 48 h post transfection, cells were infected with DENV and Cipa was added after viral adsorption. Boxplots with same labels have similar mean values and dissimilar labels have different mean values i.e both the boxplots with label “a” have same mean values while the box plots with labels (a, b, c) differ significantly with each other (P value<0.01 Anova test). Tukey test details in supplementary figure 11.

## Discussion

The use of *Cissampelos pareir*a as a hormone balancing medicine is widespread, recently the antiviral activity of the formulation also came to light. Despite this, the molecular mechanisms underlying such diverse activities of Cipa remain unexplored. We built our hypothesis around the Ayurvedic knowledge of its usage and the fact that there could be possible crosstalk between the pathways implicated in these unrelated disorders, possibly through hormone response. We carried out genome-wide expression profiling of MCF7 cell lines treated with CIPA to infer their plausible mechanisms of action as well as queried the signatures in connectivity map to infer similarities with the effect of other drugs, perturbagens, and gene-knockouts.

Interestingly, the gene set enrichment analysis highlights an enrichment corresponding to the upregulation of genes upon inhibition of ESR1 and response to endocrine therapy in cancer (Figure 1A). Upon qPCR analysis, we also find a downregulation of the ESR1 mRNA by Cipa. These results suggest that Cipa may play a role as a modulator of ESR1. When we searched for the estrogen response elements in the promoter region of the genes differentially expressed by Cipa, we found that the downregulated genes were highly enriched for such elements. Noteworthy, virtual screening of the constituent drug molecules against the estrogen receptor alpha revealed four distinct sites of binding. The possibility of multiple components binding at the same site shows inherent redundancy in the herbal extract. This could impart a synergistic activity in the whole extract of Cipa that might be lost in isolates. The analysis also revealed an abundance of the components in the formulation that have comparable binding energies to previously reported regulatory molecules.

CMAP analysis reveal that Cipa signature positively connects with protein synthesis inhibitors and the knockdown signature of genes are involved in translation and ribosomal protein processing that are needed for viral transcription and translation. The knockdown signature of the gene RPL7, a large ribosomal protein 7 and an ESR1 regulator ^11,12^ shows the highest positive connectivity with the Cipa gene signature. ESR1 is also known to regulate the immune response and estrogen regulators have been shown to possess antiviral activities ^13–16^. Protein translation inhibition is a very well established first-line defense against viral infections ^17,18^. Recently, many translation inhibitors with antiviral activity and antivirals as translation inhibitors have been reported ^19,20^. All kinds of viruses have been seen to involve the endoplasmic reticulum and host translational machinery to form a membrane structure in which they reside and replicate ^21–23^. Most interestingly, the biological pathways enriched for differential expression also include protein modification processes in upregulated and endoplasmic reticulum protein processing in downregulated genes. Cipa appears to inhibit translation without inducing cell death or inducing autophagy, hence could be potentially an effective antiviral agent.

Most striking, an HCV inhibitor Telaprevir has recently been observed to inhibit estrogen dependent proliferation of breast cancer cells by modulating levels and transcriptional activity of ERα ^24^. Also, Emetine which is the highest scoring small compound in the CMAP, has been shown to decrease the levels of ER alpha in MCF7 cells ^25^.

Our results so far indicate an involvement of the estrogen axis in the antiviral activity of Cipa. Upon experimental validation we observed that Cipa induced reduction of viral titers was markedly lower in MDA-MB-231 cells that are naturally deficient in estrogen receptor alpha and also in MCF7 cells when ESR1 gene is knocked down. Thus, signifying the effect of the estrogen axis in the antiviral action.

It is interesting to note that using a formulation with known and widespread usage, we were able to identify ESR1 as a factor involved in the antiviral activity of a therapeutic. This gives a hint of probable involvement of such connections in other viral infections as well, thus somewhat signifying a potential difference in the antiviral response between males and females.

## Materials and methods

### Cell culture and Cipa treatment

*Cissampelos pareira* L. whole plant extract was obtained commercially in lyophilized form, from a GMP certified manufacturer. Crude water extracts were reconstructed from powdered Cipa. Fresh extracts were prepared right before each experiment. The chemical profiling of the extract was done using UPLC. The details are described in supplementary methods and the identified constituents are provided in supplementary table S2.

MCF7 (triple positive breast cancer) cell line was obtained from NCCS, Pune. It was maintained in DMEM high glucose media, supplemented with 10% FBS,1M HEPES and 1X antibiotic antimycotics. The cells were kept at 37°C and 5% CO2. MycoAlert (Cat no. LT07-318) was used to test the culture for mycoplasma.

MCF7 cells were treated with 1µg, 10µg, 100µg, 500µg and 1000µg per ml of the aqueous extract from *Cissampelos pareira* L. The extract was prepared outside the cell culture hood and then filtered using 0.22µm syringe filter before its use in cell culture. The MCF7 cells were seeded at a confluency of 60-70% 18-24 hrs. prior to treatment. The cells were treated with Cipa for 24 hours for each experiment. A fresh extract was made for each set of experiments.

### RNA isolation and Whole transcriptome analysis

MCF-7 cells were seeded in 6 well plates at 60-65% confluency and then treated with vehicle, 1µg, 10µg, 100μg, 500µg and 1000µg of Cipa for 24 hours. The cells were then trypsinized and washed with PBS. Total RNA was isolated using TRIzol (Ambion, Cat no. 15596026) extraction method and its integrity was checked on 1% agarose gel which was followed by Nanodrop quantification (ND1000, Nanodrop technologies, USA). Genome-wide transcriptome data was generated using 250µg of the total RNA for each sample following the manufacturer’s protocol. GeneChip™ (Affymetrix) Human Transcriptome Array 2.0 cartridges were used for this experiment. HTA 2.0 chip can capture 245,349 protein coding transcripts and 40,914 non protein coding transcripts in a 64-format. Briefly, the total RNA was prepared with poly A controls and first strand cDNA followed by second strand cDNA were synthesized. This was followed by in vitro transcription to form cRNA which was used as a template for the formation of single strand cDNA or ss-cDNA. Finally, the ss-cDNA was fragmented and labelled. At each step, purification and quantification of the samples were ensured. Approximately 200µl of the sample, containing about 5.2µg of the labelled ss-cDNA was loaded into the cartridges, which were then kept in the Affymetrix® Hybridization Oven, set at 45°C and RPM 60, overnight. The cartridges were registered on AGCC (Affymetrix® GeneChip® Command Console) and the fluidics of the experiment (washing and staining) was done using Affymetrix® GeneChip® Fluidics Station 450. The stained chips were then scanned using GeneChip™ Scanner 3000. The preliminary images were quality checked and .cell files were generated.

The generated CEL files were background normalized using the RMA method. Batch effects were removed and differential gene expression analysis was done using the limma package in R. After background correction and RMA normalization, the probes that exceeded the p value<0.05 were annotated and analyzed for differential expression. Since no probe could qualify the cut off at 1µg, 10µg and 100µg, the differential gene expression was calculated using the dataset from 500µg and 1000µg treatments with the vehicle used as control.

### Functional enrichment of the differentially expressed genes

For functional analysis we used g: profiler ^26^ (https://biit.cs.ut.ee/gprofiler/gost). After correcting for FDR<5%, enrichments were analyzed for GO: Molecular Function, Cellular Compartment, Biological Processes, KEGG and Reactome.

In order to identify the specific sets of genes modulated by Cipa, we used Gene set enrichment analysis. We carried out a pre-ranked analysis for GSEA (Gene set enrichment analysis), in which the gene list was ordered from the highest positive gene expression to the lowest negative gene expression. The latest versions of gene sets databases were selected for query. Datasets with more than 500 and less than 15 genes were excluded from the query. The output was set to a minimum cut off of p value<0.05 and corrected for FDR<25%. An additional filter of FDR<5% was applied for identification of significant enrichments.

### Estrogen Response Elements (ERE) analysis

Sequence of 5KB upstream region of genes were downloaded from UCSC genome browser for Human genome GRCh38 build. ERE sites were mapped to 5KB upstream sequence using promo tool ^27^ with similarity cut off value of 8.7. High ERE density in 5KB upstream leads to down regulation of genes as per regression analysis. ERE density’s coefficient of slope of regressed line (β = -0.0044) is negative and p-value is significant (0.018). Pair wise ERE density comparison of differentially expressed genes and unchanged genes was done using R software (version 3.2).

### Virtual screening of constituent compounds for estrogen receptor binding affinity

Virtual screening of CIPA ligands was performed against the estrogen receptor crystal structure (PDBID: 3OS8) using Autodock Vina, a more accurate and faster version of Autodock 4. The CIPA ligands available in the PubChem database were downloaded from there, the 3D structures of the rest of the ligands were drawn using Marvin Sketch, a computational tool for drawing 3 and 2 dimensional chemical structures. The structures were randomized and minimized prior to docking. A blind docking study was performed for each ligand wherein the possible search-space was the complete receptor molecule. For each drug molecules, the docking parameters were as follows: center_x: 9.43, center_y: 22.811, center_z: 23.418, size_x: 60, size_y: 60, size_z: 60, exhaustiveness 5000, num_modes 50000, energy_range: 20. Furthermore, for each ligand, 50 such runs were performed and the conformation with the minimum binding energy from among all the stable conformations of the 50 runs (a total of 1000 conformational possibilities) was selected for the cluster analysis.

The analysis was performed using ADT, UCSF Chimera and LigPlot+.

### Connectivity map analysis for identifying similarities with known perturbations

The connectivity map dataset currently contains 1,319,138 gene expression profiles, resulting in 473,647 signatures, generated against approximately 27,000 perturbations in 9 cell lines including MCF7, HEPG2, A549, A375, PC3, HCC515, HT29, HA1E and VCAP. The differentially expressed genes were ranked according to fold change and a signature of up and downregulated genes was generated. The gene signature containing genes which were valid i.e., having a valid HGNC symbol or Entrez ID and were also present in the LINCS gene space (represented in the L1000 data as landmark or well inferred) was used to query the connectivity map using clue.io touchstone database.

### Cipa treatment in DENV infection

MCF-7 were seeded at 100,000 cells per well in 24 well plate and were maintained for 24 hours (37 degrees; 5% CO2). Virus challenge (at 10 MOI, see supplementary) was given for 1 hour, here the media was supplemented with 2% FBS. After 1-hour of viral adsorption, cells were washed with PBS and DMEM high glucose with 10% FBS with or without Cipa (50μg and 100μg) was added. DENV titers in the supernatant were determined by plaque assay at 24 h pi. DENV plaque assay was set up in BHK-21 cells. 50,000 BHK-21 cells were plated per well in 24-well plate. Serial dilutions of virus were added in triplicates and allowed to adsorb for 1 h, followed by overlay with 0.5% carboxymethyl-cellulose (CMC; Sigma). After 72 h, cells were fixed in 3.7% formalin and plaques were visualized by staining with crystal violet.

### siRNA-mediated knockdown of ESR1 in MCF7 cells

1 μM concentration of siRNA targeting ESR1 gene and non-targeting control (NTC) were mixed with Opti-MEM (Life Technologies) and 1 μl of Lipofectamine RNAiMax to a total volume of 100 μl in a 24-well plate. Cells were trypsinized and volume made up so as to contain 60,000 cells in 400 μl antibiotic-free medium. After 20 min incubation of the transfection complex, cell suspension was added into each well. Knockdown efficiency was determined by qRT-PCR at 48 h post transfection. Cells were infected with DENV-2 at 48 h post transfection. Cipa was added at a concentration of 50 µg after 1 h of viral adsorption. DENV-2 titers were measured by plaque assay at 24 h pi.

## Declarations

### Authors’ Contributions

This study was conceived by MM and BP and designed by MM, BP, MH and DD. The experiments were performed by MH, AP, AG, KS, DS, S.A.B. The Connectivity map analysis and interpretations were done by MH, DD, BP, MM and the functional validations were performed by MH, DS, and KS. The molecular docking analysis was done by PG, and lead by AR. The dengue infection experiments were performed by AP, AG, under the supervision of G.R.M. MH, DD, BP, and MM wrote the whole manuscript. All authors read and approved the final manuscript.

## Acknowledgments

The authors would like to thank Mr. Manish Kumar for imaging facility, Dr. TN Vivek for critical inputs, Himanshi Tanwar, Ratika Sehgal and Shivam Singh for technical support. The authors would like to thank Vineet Jha from Persistent Systems ltd for ERE analysis. Research fellowship support to MH (University Grants Commission), and DD (Department of Biotechnology) is duly acknowledged.

## Competing Interest

The authors declare they have no competing interests.

## Funding

1. Council of Scientific Research (CSIR) TRISUTRA (MLP-901)
2. Center of Excellence on Applied Developments in Ayurveda, Prakriti and Genomics, grant by Ministry of AYUSH (GAP0183), Govt. of India

## Notes

### Competing Interest Statement

The authors have declared no competing interest.

### Summary of Updates

The title, abstract, and introduction have been updated. The author's list has been updated. Two of the previous sections and two corresponding figures have been removed from results and material and methods.

