## Supplementary information for "Transcriptome analysis and connectivity mapping of *Cissampelos pareira* L. provides molecular links of ESR1 modulation to viral inhibition"

**Supplementary material**

**Materials and methods:**

**Cell viability assay:**

Toxicity of the extract at the aforementioned doses was done using MTT (HiMedia, Cat# TC191) assay. MCF7 cells were seeded in 96 well plates and the extracts were added. The treated cells were kept at 37˚C and 5% $CO_2$ for 24 hours. The media was then removed and MTT at a final concentration of 20µg per well was added. The cells were then kept in the incubator for 4 hours at 37˚C and 5% $CO_2$. The reduced formazan crystals were then dissolved in 100µl DMSO and kept at room temperature in the dark for 15 minutes. The absorbance was measure at 560nm using Tecan plate reader Infinite200 Pro. Each concentration was analyzed in triplicates and the experiment was repeated three times. For this assay paclitaxel at a concentration of 10µg was used as positive control.

**Phosphorylation of EIF2α by Cipa:**

To check the ratio of pEIF2α: EIF2α first in cell western was done using antibodies against pEIF2α (ProSci, Cat no. 79-226) and EIF2α (Abbkine, Cat no. ABP51245). Secondary antibodies from LI-COR, Anti-Rabbit 680 (Cat no.926-68023), Anti-mouse 680 (Cat no.926-68022) and anti-rabbit 800 (Cat no. 926-32211) were used. The MCF7 cells were seeded in a 96 well plate and treated with the extract for 24 hours. The cells were washed with PBS and fixed using 3.7% formaldehyde. After permeabilizing the cells with 0.1% triton X, blocking of the proteins was done using 3% BSA for 1.5 hours. The cells were incubated with primary antibodies overnight at 4 degrees and washed with 0.01% tween 20, followed by incubation with secondary antibodies for 2 hours. The plate was read in LI-COR Odyssey Infrared reader after thorough washing and drying. The infrared fluorescence was measured by manual selection of the wells. The values were noted and the ratio was calculated.

**HPG global protein synthesis assay:**

To assess the effect of the extract on global protein synthesis, Click-iT™ HPG Alexa Fluor™ 488 Protein Synthesis Assay (Cat no. C10428) was done according to the manufacturer's protocol. Briefly, Cells were allowed to grow overnight and the media was changed to DMEM high glucose without cysteine and methionine containing HPG. The cells were co incubated with 500µg and 1000µg extract for 3 hrs. As a positive control for protein synthesis inhibition, cycloheximide was used at the concentration of 10µg. The cells were washed with PBS, fixed with formaldehyde and permeabilized using triton-X. Substrate to HPG was added and the cells were incubated for 45 mins, counterstained with DAPI and the fluorescence was measured using High content imaging

**Collection of plant material and preparation of extract:**

Whole plant of C. pareira was collected from the Palampur, HP, India (alt. 1350 m). The identification of the plant material was done by a taxonomy expert in CSIR-IHBT, Palampur and a voucher specimen (no. PLP16688) was deposited in the herbarium of CSIR-IHBT, Palampur, HP-176,061, India.

Chemicals and reagents:

For UPLC analysis, formic acid was obtained from S. D. Fine Chemicals Ltd. (Mumbai, India) whereas methanol and water (LC grade) were purchased from J. T. Baker (Mallinckrodt Baker Inc., St. Louis, MO, USA).

The pure molecules from roots of the plant were isolated as reported by us recently (3).

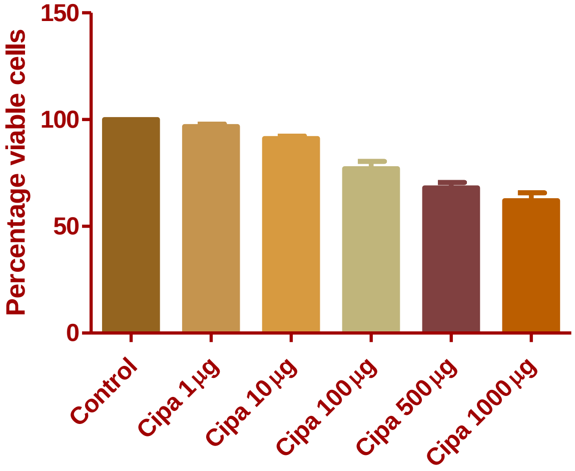


**Fig. S1.** Cipa doesn't show toxicity in MCF7 cells, except at higher concentrations. MTT cell viability assay shows less than 40% toxicity induced by Cipa even at higher concentrations.

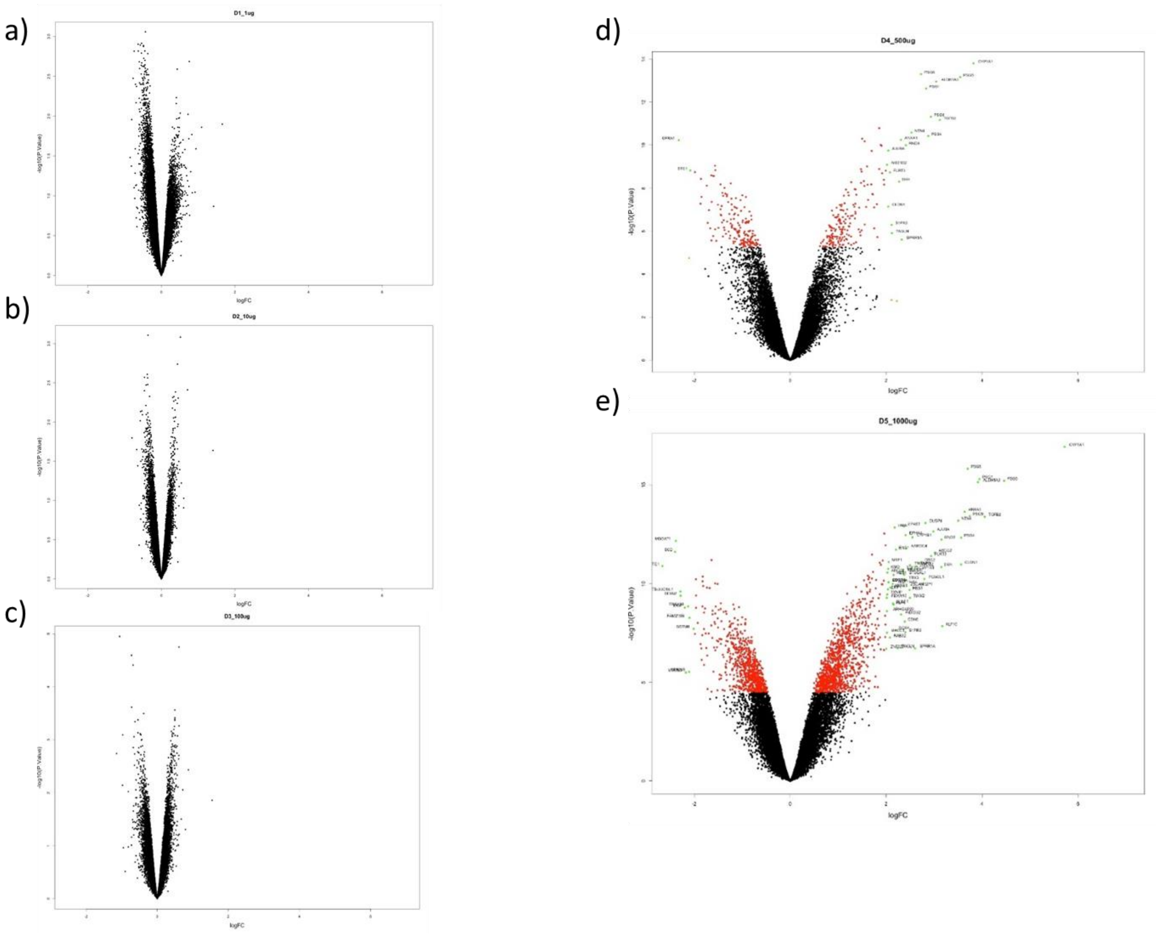


**Fig S2.** Volcano plots for probes with intensity value beyond the threshold for a-c) 1, 10 and 100μg, no probe qualifies the cut off, d) 500μg and e) 1000μg. p value<0.001, FC>|1|

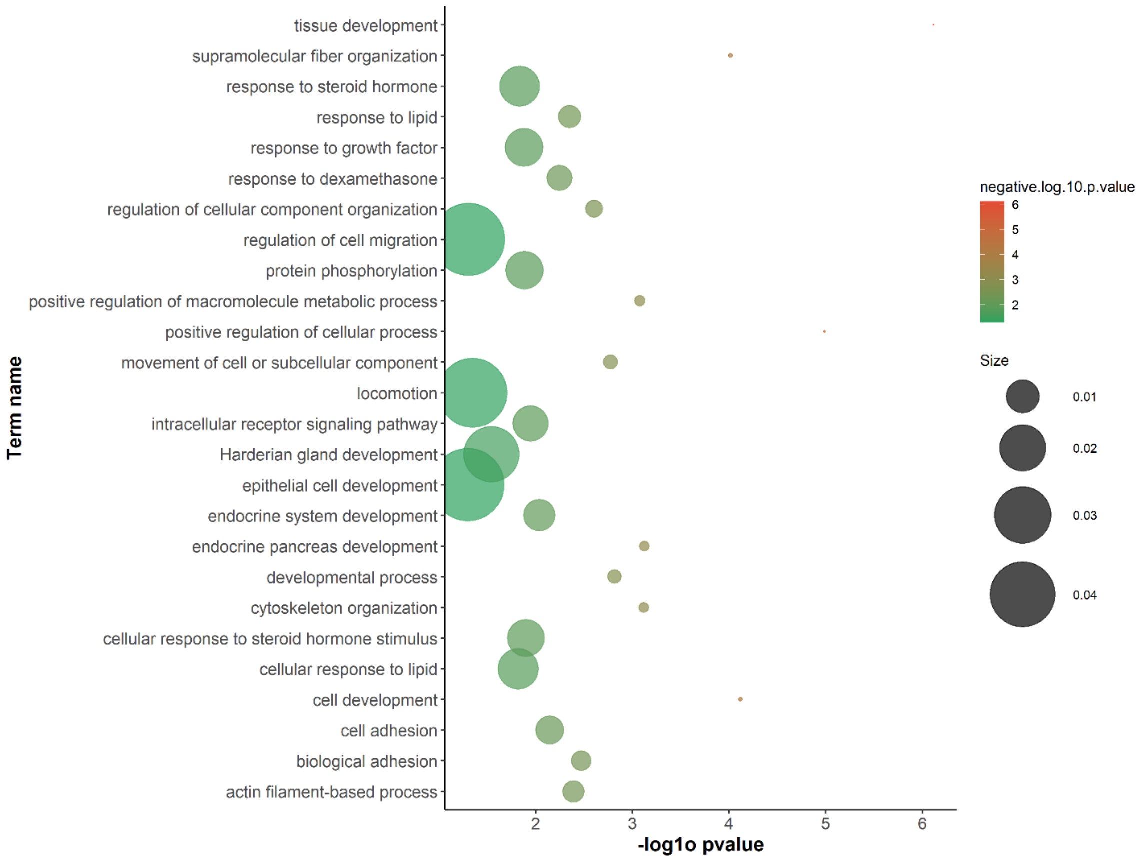


**Fig S3.** Gene ontology of the genes differentially expressed in response to Cipa using g: profiler (4). The p value was set to <0.01, FDR<0.05. This graph represents the biological processes from GO.

| Regression with ERE sites | Coefficients | P-value |
|---|---|---|
| Intercept | 0.725 | 4.756E-15 |
| ERE sites 5KB Upstream | -0.004 | 0.018 |

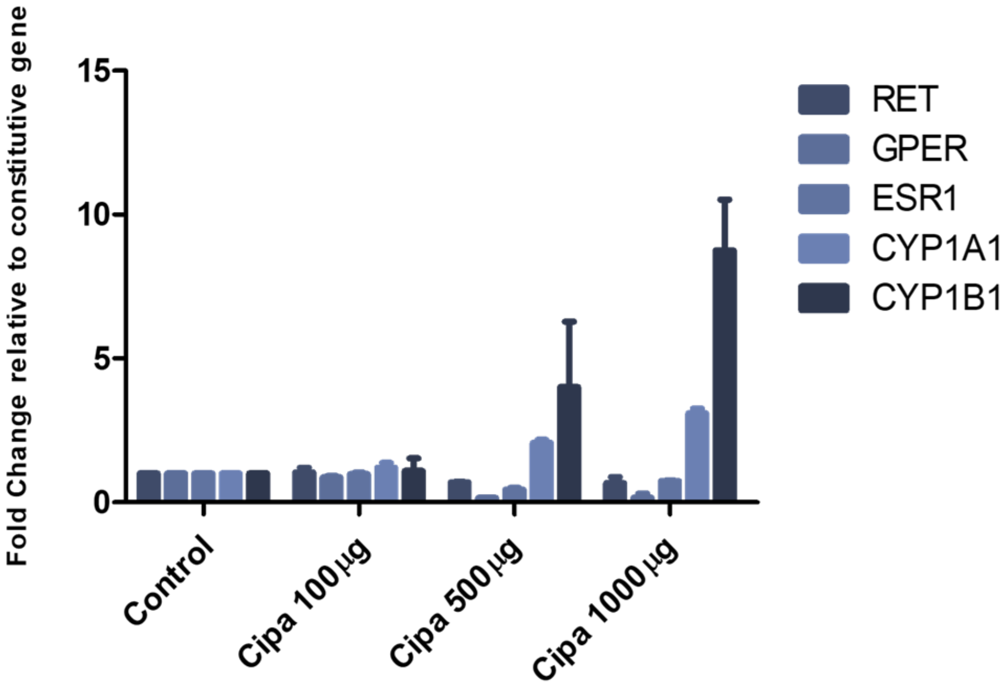


**Fig S4.** Regression values between fold change and the number of ERE sites. Response of well characterized estrogen responsive genes upon treatment of Cipa in MCF7 cells. Each of these genes have >25 ERE elements in their promoters.

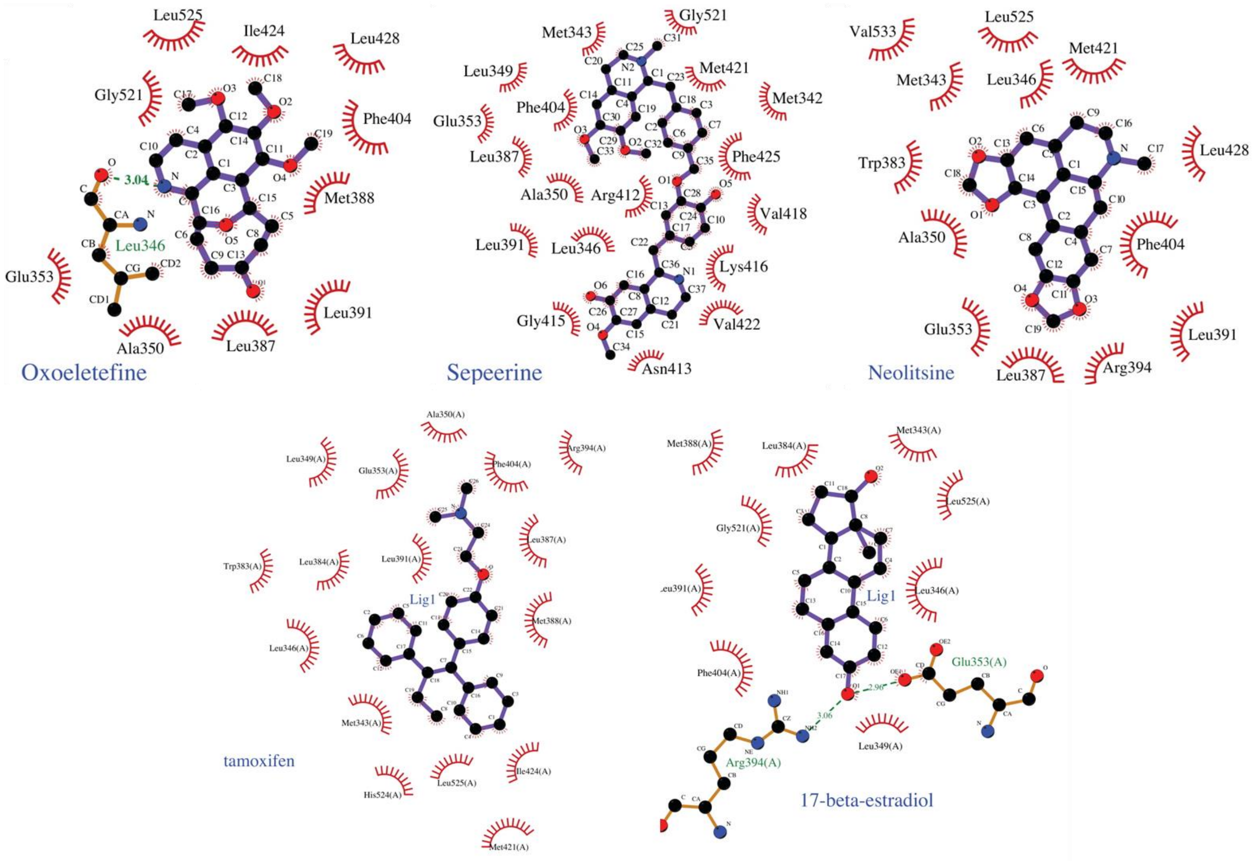


**Fig S5.** Docking details of the control molecules and the top three compounds constituent of Cipa around Erα.

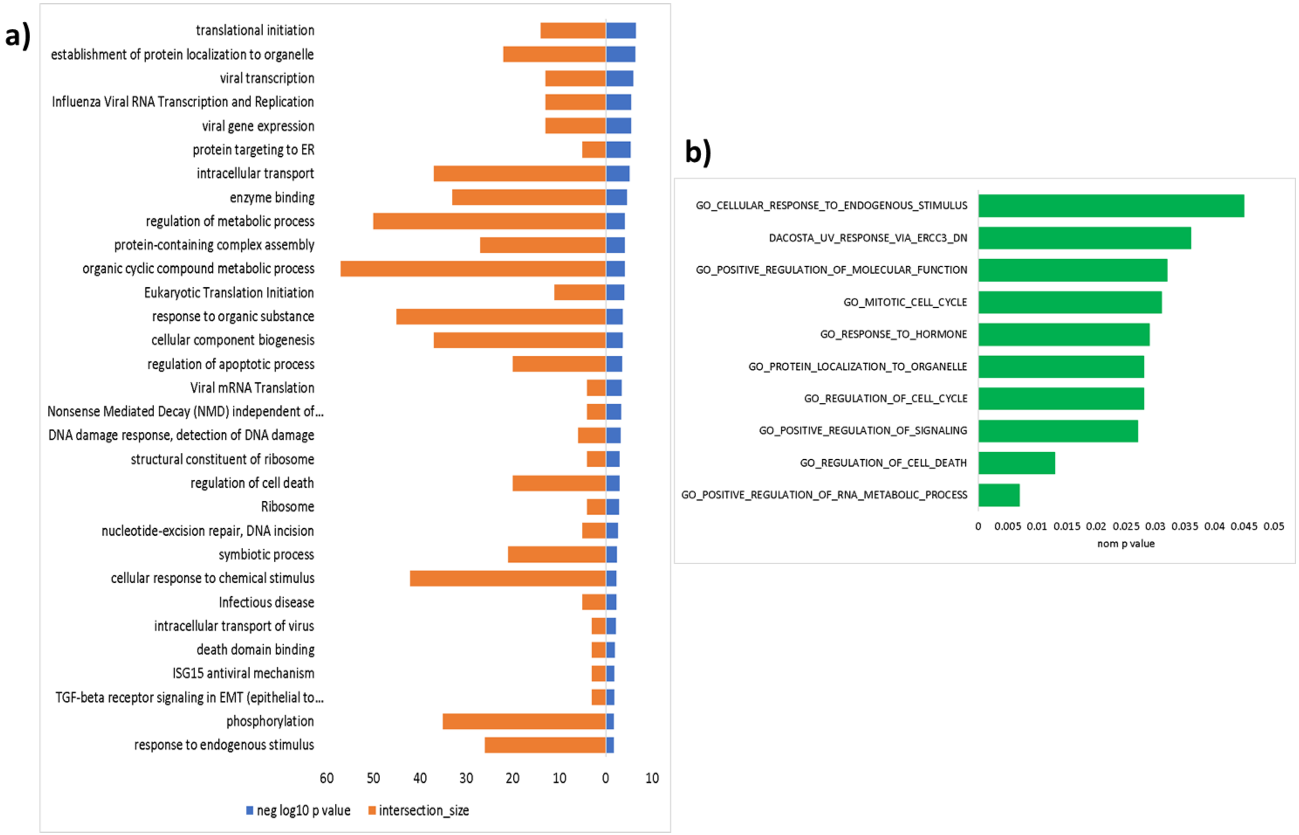


**Fig S6.** Terms enriched for genes with knockdown signature having >90 score connectivity with Cipa within CMAP (a) Biological pathways and processes and the number of genes intersecting with the total genes in the query (b) Gene sets enriched after GSEA.

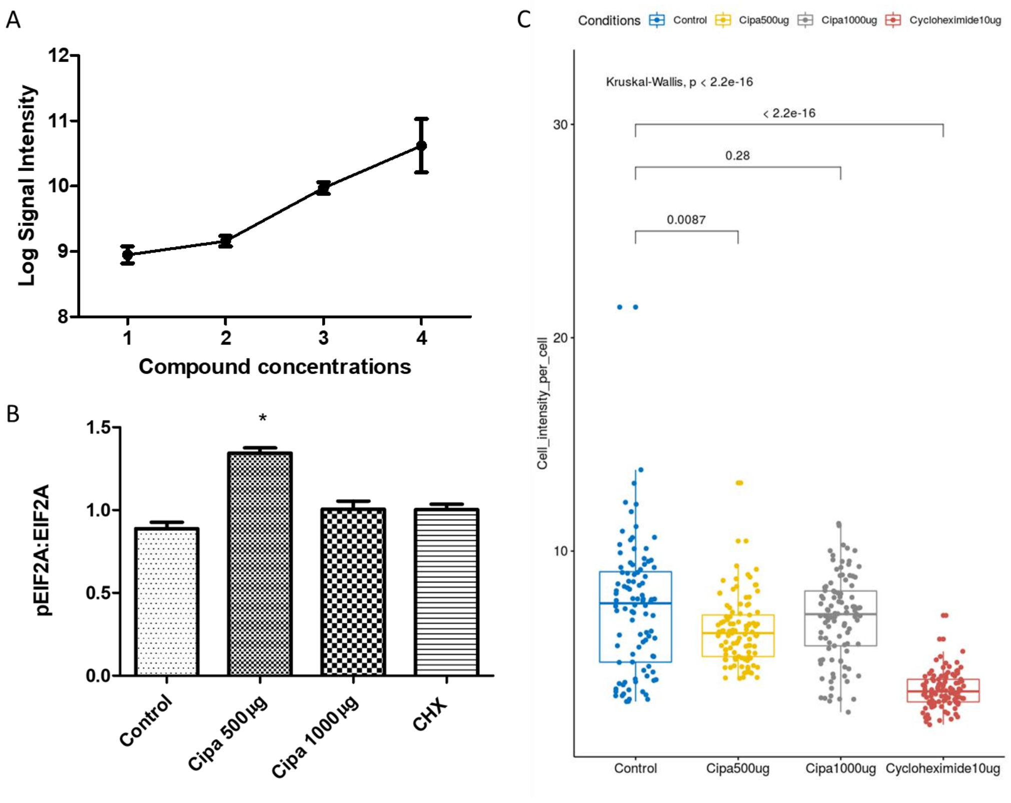


**Fig S7.** Cipa shows moderate protein translation inhibition: (A) EIF2AK3 microarray intensity values at the Cipa concentrations of 0, 100, 500 and 1000µg. (B) Ratio of phosphorylated EIF2A to non-phosphorylated EIF2A as determined by ICW. * p value=0.05 (C) Global protein translation inhibition determined using HPG assay. Fluorescent HPG replaces methionine during the formation of new proteins. The fluorescent values were measured at >100 different grid locations as intensity per grid and divided by the number of cells counted in the grid using high content imaging.

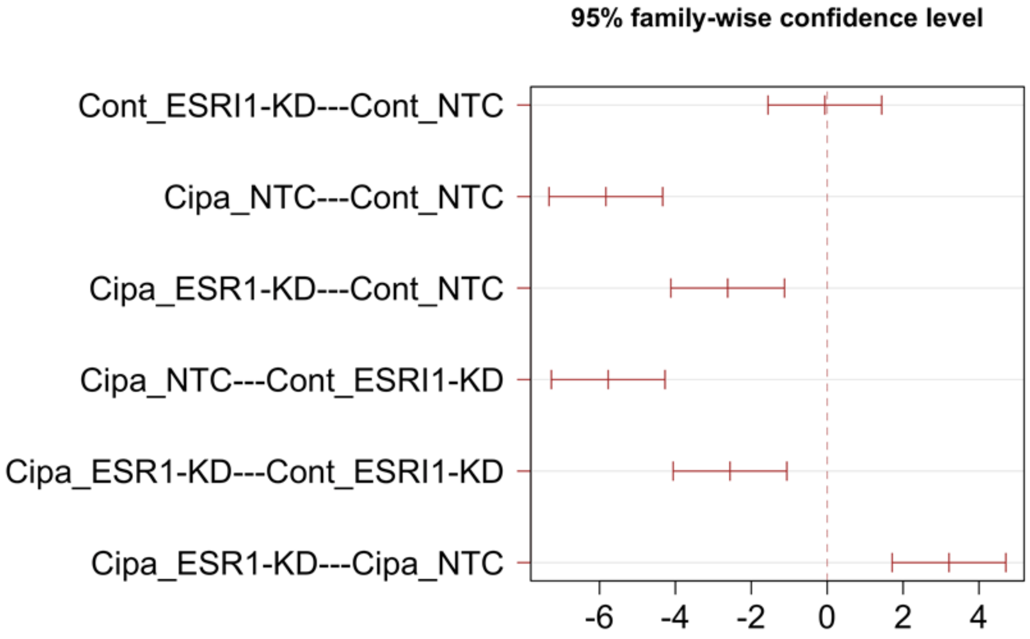


**Fig S8.** Difference in mean levels of viral titers. Tukey test compares the mean of every possible combination. Negative value indicates decrease in titers whereas positive value indicates that the mean (viral titer) is greater. Therefore, for conditions 1-5, there is an inhibition in the viral titers, however for the 6$^{th}$ condition, the inhibition is less compared to Cipa_NTC.

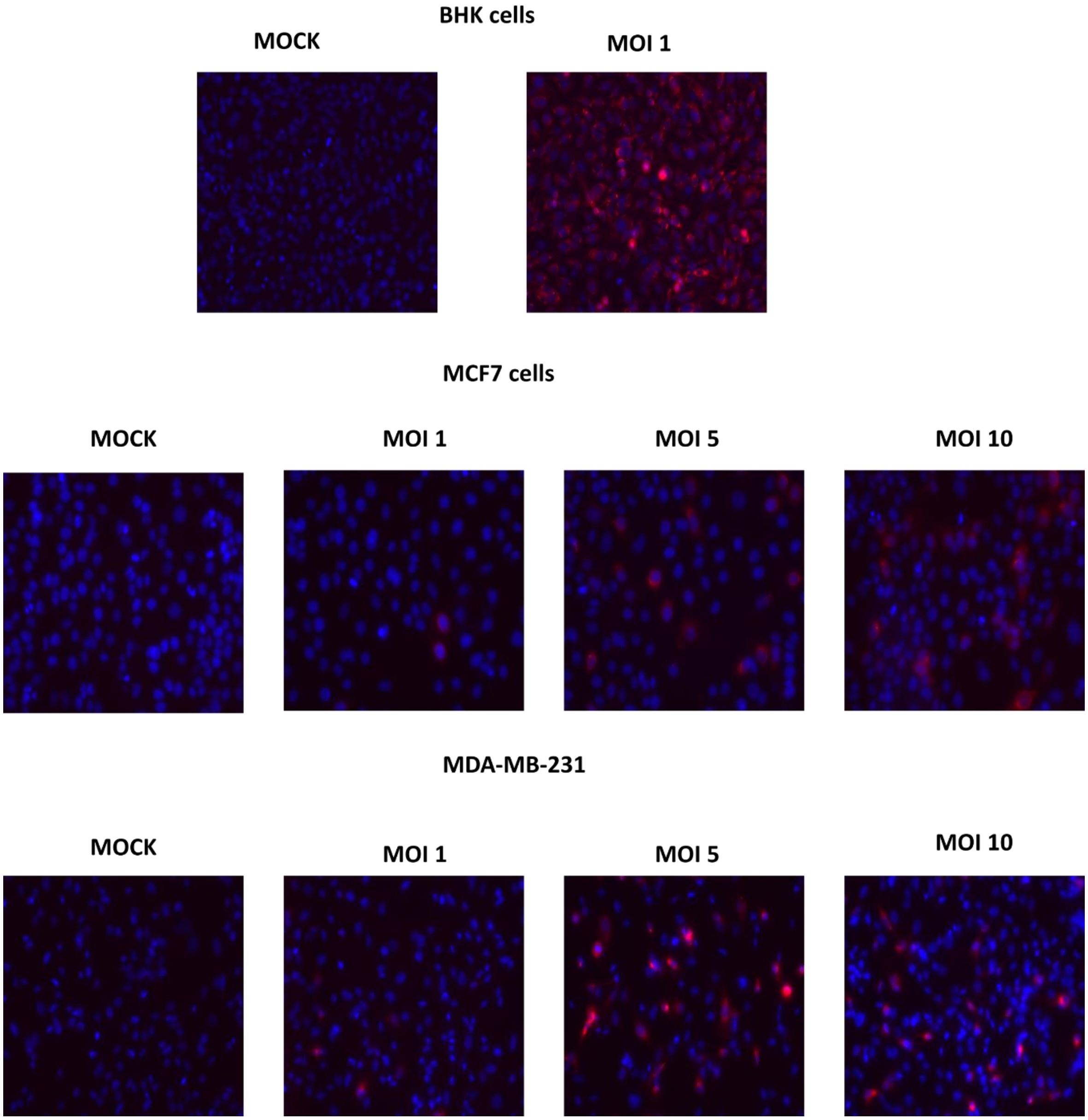


**Fig S9.** MOI of dengue virus in MCF7 and MDA-MB-231, the blue stains indicate the nuclei of the cells (DAPI) and the red stains indicate the viral proteins.

**Table S1. Cipa constituent compounds, their minimum and maximum energies with ER-alpha, the binding cluster each fall into.**

| DRUG NAME | Minimum Binding Energy(kcal/mole) | Maximum Binding Energy (kcal/mole) | Cluster |
|---|---|---|---|
| Sepeerine | -10.2 | -5.8 | |
| 17-beta-estradiol (Control) | -10.1 | | 1 |
| Neolitsine | -9.6 | -5.5 | 1 |
| Oxoeletefine | -9.6 | -4.7 | 1 |
| Cycleanine | -9.5 | -7.5 | 1 |
| Liriodenine | -9.5 | -5.9 | 1 |
| Pareitropone | -9.4 | -5.2 | 1 |
| Tamoxifen (Control) | -9.2 | | 1 |
| Pronuciferine | -9.1 | -4.8 | 1 |
| Bulbocapnine | -9 | -5.4 | 1 |
| Cissaglaberrimine | -9 | -5.7 | 1 |
| Grandirubrine | -8.8 | -4.9 | 1 |
| Quercetin | -8.8 | -5.4 | 1 |
| Eletefine | -8.7 | -4.7 | 1 |
| Norruffscine | -8.7 | -5.2 | 1 |
| Pareirubrine_B | -8.7 | -5.1 | 1 |
| Oxobuxifoline | -8.6 | -5.7 | 1 |
| Trilobinine | -8.6 | -5.2 | 1 |
| Cissacapine | -8.6 | -6.9 | |
| Arachidic_acid | -8.5 | -5.7 | 1 |
| Cissamine | -8.4 | -5.1 | 1 |
| Magnoflorine | -8.4 | -5.2 | 1 |
| Norimeluteine | -8.4 | -4.8 | 1 |
| Methylwarifteine | -8.4 | -6.1 | 3 |
| Pelosine | -8.4 | -6 | |
| Corydine | -8.3 | -4.8 | 1 |
| Crotsparine | -8.3 | -4.9 | 1 |
| Glaziovine | -8.3 | -5 | 1 |
| Insulanoline | -8.3 | -6.2 | 2 |
| (S)-6-Methoxyjuziphine | -8.3 | -4.9 | |
| corytuberine | -8.2 | -5.2 | 1 |
| Cissampeloflavone | -8.2 | -5.9 | 2 |
| Nuciferine | -8.1 | -5 | 1 |
| Reticuline | -8.1 | -5 | 1 |

| DRUG NAME | Minimum Binding Energy(kcal/mole) | Maximum Binding Energy (kcal/mole) | Cluster |
|---|---|---|---|
| **Roraimine** | -8.1 | -6.1 | 3 |
| **Milonine** | -8 | -4.7 | 1 |
| **N.N-Dimethyllindcarpine** | -8 | -4.8 | 1 |
| **Cissampentin** | -8 | -6.2 | 3 |
| **Hayatidine** | -7.9 | -6.2 | 3 |
| **Dimethylwarifteine** | -7.9 | -5.7 | 4 |
| **Hayatine** | -7.8 | -6 | 3 |
| **Laurifoline** | -7.7 | -5.2 | 1 |
| **Isochondrodendrine** | -7.7 | -6.2 | 2 |
| **Berberine** | -7.6 | -5.2 | 2 |
| **Cissampareine** | -7.6 | -6.3 | 3 |
| **Dehydrodicentrine** | -7.4 | -5.4 | 1 |
| **Dicentrine** | -7.4 | -5.3 | 1 |
| **Salutaridine** | -7.4 | -4.6 | 1 |
| **Hayatinin** | -7.4 | -6 | 2 |
| **Insularine** | -7.3 | -5.8 | 2 |
| **Reserpine** | -7.3 | -5.8 | 2 |
| **Warifteine** | -7.3 | -5.4 | 4 |
| **Isoimerubrine** | -7.2 | -4.9 | 1 |
| **Laudanosine** | -7.1 | -4.7 | 1 |
| **Deca-2E.4E-dienoic_acid_isobutylamide** | -6.7 | -4 | 1 |
| **Lauroscholtzine** | -6.7 | -4.9 | 1 |
| **Pareirubrine_A** | -6.7 | -4.8 | 4 |
| **Decene-2-oic_acid_isobutylamide** | -6.4 | -3.7 | 1 |
| **Decanoic_acid_isobutylamide** | -6.1 | -3.4 | 1 |
| **Thymol** | -6.1 | -4.2 | 1 |
| **Magniflorin** | -6.1 | -4.9 | 2 |
| **Galacturonic_acid** | -5.9 | -4.3 | |
| **d-Quercitol** | -5.7 | -3.8 | |

**Table S2: Antiviral activity of positively connected compounds**

| Compound | Virus | References | Year |
|---|---|---|---|
| Emetine | Coronavirus | PMID: 24841273 | 2014 |
| | New Casle Disease, Bovine Herpervirus, Buffalopoxvirus | PMID: 28624461 | 2017 |
| | Human Cytomegalovirus | PMID: 27336364 | 2016 |
| | Rabies virus | PMID: 30028873 | 2018 |
| | HIV-1 | PMID: 26111177 | 2015 |
| | Enterovirus | PMID: 31734270 | 2020 |
| | HSV-2, HIV-1, Influenza | PMID: 31635418 | 2019 |
| | Novel Coronavirus | PMID: 30918074 | 2020 |
| | Zika and Ebola virus | PMID: 29872540 | 2018 |
| Anisomycin | Coronavirus | PMID: 24841273 | 2014 |
| | Poliovirus | PMID: 18243348 | 2008 |
| | Dengue and Zika virus | PMID: 32081740 | 2020 |
| Q-XII-47 | Dengue virus | PMID: 28034743 | 2017 |
| | Novel Coronavirus | *Preprint* | 2020 |
| Homoharringtonine | Coronavirus | PMID: 24841273 | 2014 |

| | | | |
|---|---|---|---|
| | Vesicular stomatitis virus, Newcastle disease virus | PMID: 30388805 | 2018 |
| | Coronavirus | PMID: 25451075 | 2015 |
| | Hepatitis B virus, Bovine diarrhoea virus | PMID: 17458779 | 2007 |
| | Foot and mouth disease virus | PMID: 31032977 | 2019 |
| Cycloheximide | HIV-1, influenza viruses, coxsackie B virus, EV71 | PMID: 21351439 | 2010 |
| | Coronavirus | PMID: 21430047 | 2011 |
| | Coronavirus | PMID: 24841273 | 2014 |
| | Coronavirus | PMID: 21430047 | 2011 |
| | Foot and mouth disease virus | PMID: 15649564 | 2005 |
| | Human T-lymphotropic virus | PMID: 22214262 | 2012 |
| | Adenovirus 1,2, 5 | PMID: 11562975 | 2001 |
| Cephaeline | Zika and Ebola virus | PMID: 29872540 | 2018 |

**Table S3: Amounts (mg/g) of compounds quantified in different samples of *C. pareira***

| **S. No.** | **Sample** | **Magnoflorine** (US-CP-7) **(1)** | **Pareirarine (2)** | **Salutaridine** (US-DR-CP-2) **(3)** | **Cissamine** (US-CP-3) **(4)** | **Hayatinine** (US-50) **(5)** | **Total alkaloids** |
|---|---|---|---|---|---|---|---|
| 1. | USCPWP-PE-50 | 12.9 | 5.4 | NQ | 18.6 | 9.5 | 46.4 |
| 2. | USCPR-PE-R | 3.5 | 1.6 | NQ | 6.0 | 1.8 | 12.9 |
| 3. | USCP-PE | NQ | NQ | NQ | 3.3 | NQ | 3.3 |
| 4. | USCP-PE-EF | NQ | 7.6 | NQ | 8.0 | NQ | 15.6 |
| 5. | USCP-PE-BF | NQ | 10.3 | NQ | 1.7 | NQ | 12.0 |
